# Chasing a moving target: Detection of mitochondrial heteroplasmy for clinical diagnostics

**DOI:** 10.1101/222109

**Authors:** Eric J. White, Tristen Ross, Edgardo Lopez, Anastasia Nikiforov, Christopher Gault, Rebecca Batorsky, Christopher Darcy, Dean R. Campagna, Mark D. Fleming, John F. Thompson

## Abstract

Clinical interpretation of human mitochondrial DNA (mtDNA) variants has been challenging for technical and biological reasons but the involvement of dysfunctional mitochondria in many diseases makes it imperative to have a validated assay for detecting pathogenic variants. We have tested several methods to identify those best suited to detect and confirm mtDNA variants. The choice of methods is dependent on the amount of DNA available for testing and the sensitivity required for detecting low-level heteroplasmies. There is a tradeoff between a polymerase’s ability to amplify small amounts of DNA and its ability to generate accurate sequence. We report a simple method to measure heteroplasmy levels of large deletions from NGS data alone without need for qPCR or other methods. Use of HapMap samples for standardization needs to be done with caution as most have novel heteroplasmic sites that have arisen during immortalization/cell culture processes. Different batches of DNA can have variable sequence. In contrast, we observed no *de novo* heteroplasmies in healthy mother-child pairs studied using blood or saliva though the frequency of pre-existing heteroplasmies often changed dramatically across generations. Long-read nanopore sequencing of individuals with two heteroplasmies suggested a random distribution of variants on single molecules but technical artifacts prevent certainty on this finding. Urine provides an additional readily accessible source of mtDNA that can be used for bone marrow transplant recipients whose saliva/blood mtDNA may be contaminated by the BMT donor’s mtDNA. We have characterized cells suspended in urine via expression profiling and shown them to be primarily mucosal cells that are independent of blood. Understanding the pitfalls of the various mtDNA sequencing methods allows development of reliable and accurate tests suitable for clinical diagnostics.

**Author Summary:** Mitochondrial DNA is important for many diseases but it is present at many copies per cell so is harder to check for mutations compared to nuclear DNA. We have studied mitochondrial DNA in different ways to see how it changes across generations and in different locations in the body. The tests need to be much more sensitive than nuclear DNA tests so that we can detect mutations down to 1%. We have shown that mitochondrial DNA changes when cell lines are used but saliva, blood and cells in the urine can all be used for testing. Cells in the urine originate as mucosal cells and are independent of blood. We developed a new method for analyzing large deletions that means sequencing data alone can be used for measuring the frequency of deletions. We also followed a family with two variable sites to better understand how mitochondrial DNA changes from mother to child. In some children, the variants stayed the same while, in others, variants disappeared.

## Introduction

Mitochondrial DNA (mtDNA) has been extensively studied because its unique inheritance and variant patterns have important implications for evolutionary biology, genealogy, forensics, human disease, and other fields (1–5). mtDNA is less than 0.001% the length of the remainder of the human genome, but contains 37 tightly packed genes, all of which are essential for life. The number of mitochondria per cell varies widely among different cell types (6). The unique maternal inheritance pattern and high mutation rate can lead to a continuous range of variant frequencies within cells/tissues with such variant sites referred to as heteroplasmic (7). The heteroplasmy can be inherited or occur *de novo* and its frequency can change with age as well as across tissues, making it challenging to understand how any given variant affects health and disease. Generally, a pathogenic variant needs to reach a frequency of >60% (deletions) or >80% (SNVs) in order to generate a phenotypic effect (8) although there are exceptions to this (9). Heteroplasmy levels may also change with age, an effect that has complicated attempts to correct mitochondrial mutations (10).

Metabolically-active cells like muscle, brain, and kidney have more mitochondria than other tissues and these cells are generally more affected by mitochondrial dysfunction. Determining whether mitochondrial variants are functionally relevant is more difficult than with nuclear variants due to the uncertain impact of heteroplasmy and the possibility that heteroplasmy varies across tissues. Comparisons across different tissues have shown significant variation in heteroplasmy frequency with the greatest variant accumulation seen in liver, kidney, and skeletal muscle (11, 12). Correlating heteroplasmy variation in different tissues with disease severity induced by the A3243G mutation concluded that heteroplasmy in urinary cells is the best predictor of disease (13, 14). These observations have led to the consensus recommendation that blood or tissue measurement of heteroplasmy levels be followed up by assessment in urine cells (15). Assigning disease relevance to mitochondrial mutations is even further complicated by the fact that hundreds of nuclear-encoded genes are known to affect mitochondrial function (16). Mutations in nuclear genes have the potential to cause either Primary Mitochondrial Disease (PMD) or Secondary Mitochondrial Dysfunction (SMD) (3).

Only recently has it been possible to properly assess heteroplasmy. The original methodology for detecting it, Sanger sequencing, is poorly suited for the task. Sanger is unreliable at detecting low but potentially biologically-relevant rates of heteroplasmy. As a result, many early studies set the minimal detection rate for heteroplasmies at 20% or higher. Furthermore, the difficulty of Sanger sequencing the whole mitochondrial genome led many to focus only on selected segments leading to a limited view of the potential impact of functional variants. The introduction of massively parallel NGS technologies with their digital determination of base calls has allowed both easy sequencing of whole mitochondrial genomes as well as more accurate determination of heteroplasmy down to very low levels (17). However, there are limits to even these technologies because the inherent error rates in single NGS reads and the need for amplification during and sometimes prior to sequencing can introduce technical artifacts. Superimposed on these technical issues is the biological issue that DNA fragments with high homology to mtDNA pepper the nuclear genome (NuMTs) and can lead to erroneous alignment of nuclear reads to mtDNA (18). The exact boundaries of NuMTs vary depending on how one defines the degree of identity required. They are typically identified by BLAST versus the nuclear genome and, in some cases, results are restricted to specific thresholds such as >500bp and >80% identity (19). All these issues can lead to technology-specific errors in determination of heteroplasmy levels.

To ensure that calling of mitochondrial mutations and heteroplasmy can be achieved with clinical quality, we have examined mtDNA using a variety of sequencing technologies, sample preparation methodologies, and tissue sources and developed a method for determining heteroplasmy for large deletions using only NGS data. We have examined the cells found in the urine to better understand their source and relevance. We find that careful choice of methods is required if sensitive and accurate sequencing is to be achieved.

## Results

mtDNA sequencing is challenging for many reasons, making it useful to compare independent methods in order to obtain optimal results. We have used two different DNA enrichment methods, hybridization capture and amplification, to select mtDNA for sequencing. Both hybridization capture and short-range PCR have the same disadvantage that the probes/primers cover only relatively short regions and thus are prone to capturing/amplifying homologous nuclear regions. When testing hybridization capture, we used probes developed by Agilent to capture mtDNA at the same time as the whole exome so that all nuclear coding regions relevant to mitochondrial function could be sequenced simultaneously with the mtDNA (20). The ratio of mitochondrial probes to nuclear probes was titrated to generate sufficient coverage across the whole mtDNA genome while minimizing coverage loss of nuclear exomic regions. While these capture probes could potentially pull down NuMT sequences as well as mtDNA, the high ratio of mitochondrial to nuclear DNA in cells ensures that NuMT sequences should be, at most, a minor fraction of the total sequence reads. If NuMTs were to be found, they would appear as variants with low heteroplasmy. Thus, NuMT contamination could limit the sensitivity of heteroplasmy detection when hybridization capture is used but the impact would be limited to low level heteroplasmy (<5%).

We found that different methods of extracting DNA from the same blood samples yielded different ratios of mitochondrial:nuclear DNA, thus necessitating different ratios of mitochondrial baits depending on which method was used for extraction. When manual extraction of DNA from blood with a Qiagen column was used, there was 3x as much mitochondrial coverage (~300x) as nuclear coverage (~100x) with a 1:50 mitochondrial:nuclear bait ratio. However, DNA extraction with an automated Autogen FlexSTAR yielded less mitochondrial coverage (~100x) relative to nuclear DNA (~100x). While 100x coverage would be sufficient for heteroplasmy detection of >5% if coverage were perfectly even, the variation in coverage across the mitochondrial genome meant that there would be limited sensitivity in some regions. As a result, bait coverage for samples prepared in an automated manner was adjusted to 1:10, which produced sufficient mitochondrial coverage (~300x mito vs. ~100x exomic) for sensitive heteroplasmy detection.

While it is desirable to obtain samples from affected tissues, it is not always possible as tissues like brain and heart are generally inaccessible to sampling. Saliva, buccal swabs, and blood are more readily obtained but may not always provide independent information as it has been found that a substantial amount of DNA in saliva/buccal swabs can originate from blood cells. When buccal DNA from individuals who have received bone marrow transplants (BMT) was examined, typically 10–30% (21) of the DNA was from the BMT donor’s blood and, on occasion, up to 100% of the DNA could be from the BMT donor. Thus, sequence determined from buccal or saliva cells may be uninformative for understanding mtDNA variants. Another readily accessible source of cellular DNA is urine where cells can accumulate from different sources. Using qPCR, we compared the relative amounts of mtDNA/nuclear DNA in selected tissues. As shown in Table 1, the amount of mtDNA in urine cells is far higher than found in blood or saliva though less than that found in liver, muscle, and kidney.

**Table 1:** Ratio of mtDNA to nuclear DNA in different tissues

|  | Blood | Saliva | Buccal | Cells in Urine | Cell-free Urine | Immortalized Cells | Liver | Muscle | Kidney Cortex | Kidney Medulla |
| --- | --- | --- | --- | --- | --- | --- | --- | --- | --- | --- |
| Sample 1 | 67 | 104 | 221 | 2323 | 569 |  |  |  |  |  |
| Sample 2 | 92 | 109 | 151 | 448 | 182 |  |  |  |  |  |
| Other |  |  |  |  |  | 255 | 3350 | 7231 | 3350 | 1710 |

When amplification is used for mtDNA selection rather than bait-based capture, care must be taken to ensure proper primer design. Primers have been designed to generate short amplicons so that even degraded DNA could be sequenced (22), up through very long amplicons that require fully intact mtDNA (23). We have evaluated amplification using both short (<500bp) and long (>8kb) amplicons to compare performance.

HapMap DNA NA12878 was amplified with short amplicons using the Ion AmpliSeq Mitochondrial panel kit followed by sequencing with the Ion Torrent PGM (22). In addition to the risk that short-amplicons could permit NuMT contamination, it was found that sequencing uniformity was poor. In a sample where median coverage was 641x, more than 5% of the mtDNA was covered at less than 20x. At the other end of the spectrum, the top 5% of the genome was covered at >35,000x and more than 2% was covered at >100,000x. These coverage extremes make it difficult to get high quality sequence over all regions, particularly if low level heteroplasmy detection is desired. Such extremes may be permissible if complete coverage is not necessary as is the case in many forensic and genealogical applications, but it is not acceptable if one needs to detect all rare variants as in clinical applications. For this reason, short-amplicon PCR was not explored further for clinical diagnostics.

Long-range PCR is advantageous for avoiding NuMT contamination and achieving more uniform coverage but makes it more challenging to obtain robust amplification. Primers need to be designed so as to be specific for mtDNA rather than NuMTs and they must also avoid common variants to prevent unequal amplification. There are only one or two NuMTs >6000 bp (18, 19), so appropriately designed longer amplicons can avoid amplifying these potential artifacts. Because of the high mutation rate in mtDNA (>400 variant sites having population frequencies >1%), it is challenging to avoid all such positions while still maximizing mismatches with NuMTs. Two sets of four primers (Table 2) were identified that meet these criteria and these primer sets amplify the entire mtDNA with just two amplicons each. This minimizes coverage variation and eliminates the chance of inadvertently amplifying known NuMTs. By using two independent primer sets, the chance of allele dropout due to SNVs or deletions is minimized.

**Table 2:**
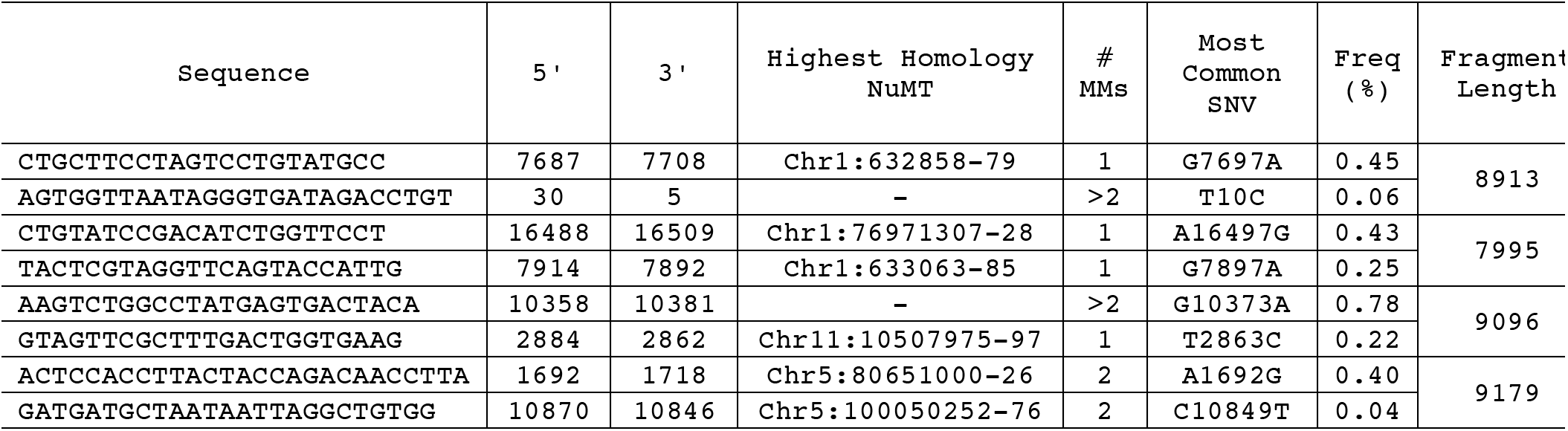
Long-range PCR primers

Of the eight primers we have validated, one is identical and one is highly overlapping with previously published primers (24). Two of the eight primers have only a single change relative to reference NuMTs while all others have 2–5 nucleotide changes from the most homologous NuMT region. None of the primer pairs should bind within a single NuMT. The most common SNV within each primer site is listed in Table 2 and no primer overlaps a SNV that has a frequency of >1% based on Mitomap.org. This combination of features provides robust and specific amplification for all samples tested and from a variety of tissue sources (Fig 1). The resultant DNA is readily sequenced via Sanger or NGS methods.

**Fig 1.**
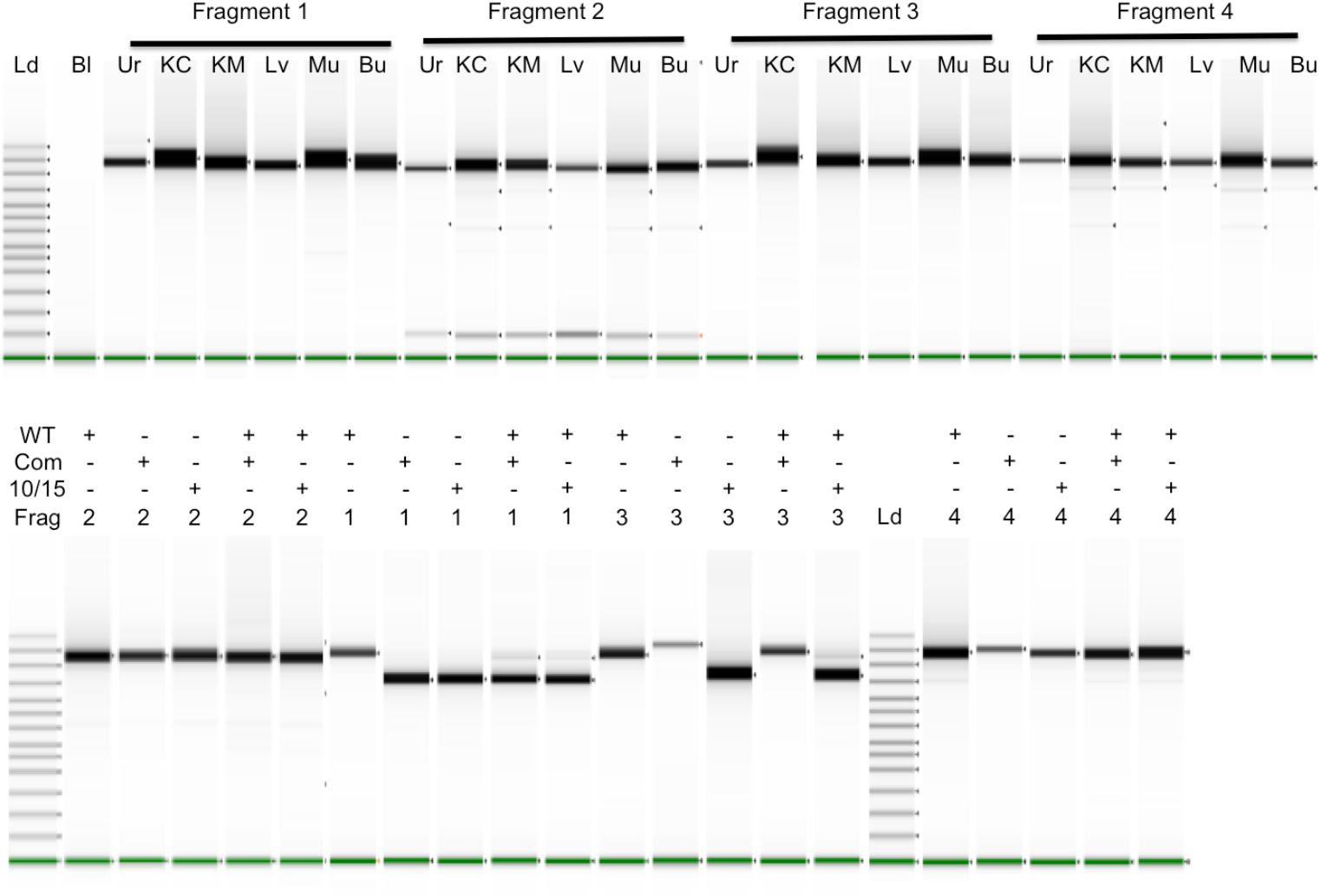
Long-range PCR products. Long-range PCR fragments were amplified individually and run on a TapeStation. In the top panel, six different tissues were amplified for each fragment: Urine cells (Ur), Kidney Cortex (KC), Kidney Medulla (KM), Liver (Lv), Muscle (Mu), and Buccal swabs (Bu). The top bands in the ladder lane on the left are 48,500, 15,000, 7000, and 4000 bp long. The Bl (blank) lane contains no input DNA. In the lower panel, mtDNA of normal length (WT) and with deletions of 5025 bp (10/15) and 4977 bp (Com) were run alone or together as indicated in each well. mtDNAs with deletions also contained heteroplasmic normal length DNA. With both experiments shown below, lanes from multiple runs have been spliced together.

In addition to the challenges created by mtDNA/nuDNA sequences, there is also the challenge of sequence quality when the need for extensive amplification is combined with the desire to measure heteroplasmy down to levels of 1% or less. As the limits for detection are pushed lower, the possibility of amplification and/or sequencing artifacts increases. As shown in Fig 2, the long range PCR input DNA requirements for different enzymes are not the same. We have found that Q5 requires more input DNA than Takara LA. Takara LA generally works with as little as 1 ng input DNA while Q5 requires at least 50 ng. Tissues such as blood and saliva yield ample DNA and allow optimal amounts of DNA for whatever enzyme is chosen. However, other tissues such as urine yield very limited DNA quantities and 50 ng can be challenging to obtain when attempting to amplify those samples.

**Fig 2.**
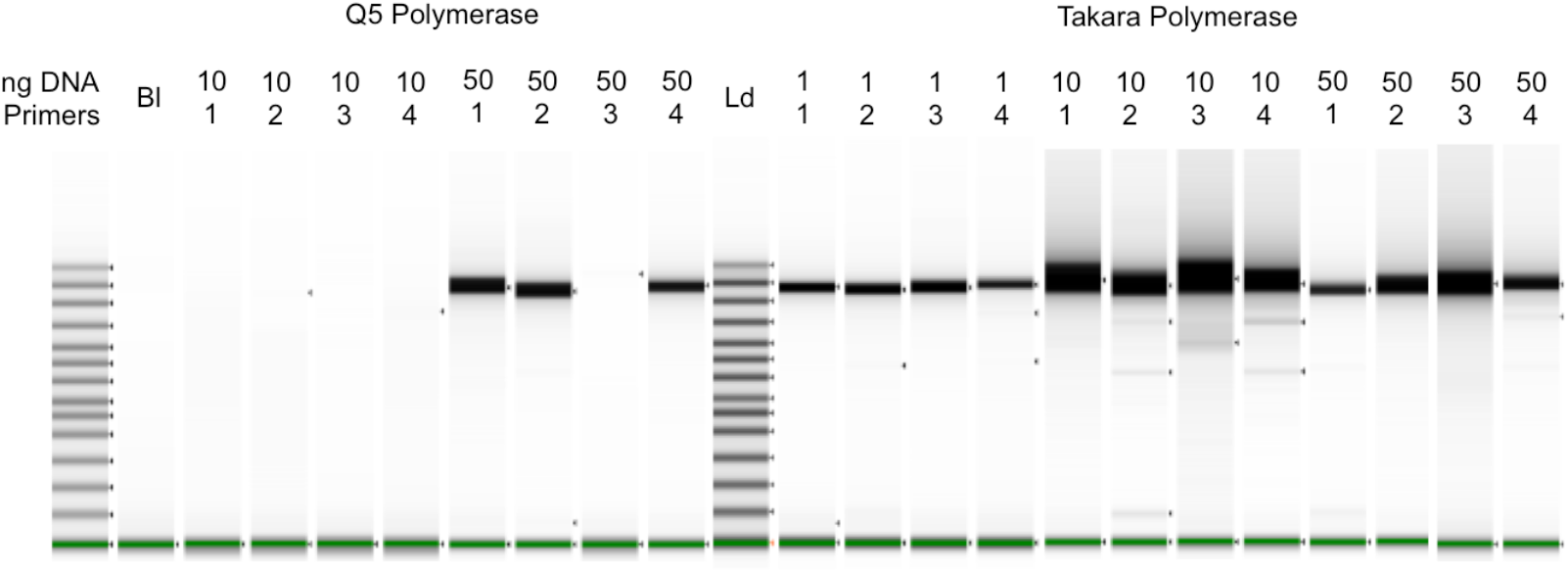
Titration of Input DNA with different DNA polymerases. Long-range PCR fragments were amplified individually and run on a TapeStation. The primers listed in Table 2 were used with varying amounts of input DNA (1–50ng) and either Q5 or Takara LA polymerase. Lanes were not normalized for intensity. Lanes from multiple runs have been spliced together in this figure.

While effective amplification is obviously a prerequisite for sequencing, it is also important to consider how the different polymerases cause sequence errors that could be significant at low heteroplasmy levels. While lower amounts of DNA can be used with Takara LA, it lacks proofreading activity so is more error-prone than Q5 (25, 26). This can lead to a higher error baseline that could be mistaken for low-level heteroplasmy. To test this, 50 ng samples were amplified with the two enzymes and sequencing results compared. When a 5% heteroplasmy cutoff is used, the results with the enzymes are identical. However, if one examines the 1–5% variant range, there are more putative low heteroplasmy variants with Takara than Q5 (typically about 6–7 with Takara and 1–2 with Q5). These variants generally differ from each other but the same false positives often occur in different samples prepared together. Thus, choice of the most appropriate enzyme is a trade-off between amplification efficiency and amplification-induced errors that can interfere with low-level heteroplasmy determinations.

In addition to pathogenic single nucleotide variants, mtDNA is also subject to large, pathogenic deletions. While these deletions can have a wide range of sizes, there is a 13 bp direct repeat that often leads to the deletion of 4977 bp between 8483–13459 to yield what is known as the common deletion (27, 28). We have included such a sample as well as another containing a deletion of 5026 bp between 10381–15406 (which we refer to as 10/15). Cells containing these deletions would not be viable without some full-length mtDNA in a heteroplasmic state.

Long range PCR of the deletion samples is shown in the lower panel of Fig 1 where a normal length mtDNA is compared to mtDNA with deletions and with mixtures of deleted/full-length DNAs. Because amplification of shorter fragments is more efficient, deletions can alter the apparent frequency of the deleted segment. Fragment 2 does not include the deletion (Supplementary Fig 1) so amplification of the pure and mixed DNAs yields identical products. Fragment 1 spans both deletions so the full-length DNA yields the expected length fragment while both deletions yield primarily the shorter, deleted fragment with low levels of full-length fragment. When the deleted DNA is combined with another full-length mtDNA in equal amounts prior to amplification, the shorter fragment still predominates with the larger fragment much lower than would be expected for an equimolar mixture. One primer for Fragment 4 (10870–10846) is within both deleted regions so cannot be used for examining the deletions because all amplified material corresponds to the full length DNA. Fragment 3 behaves differently with the two deletions because one primer (10358–10381) is missing from the common deletion region while still present in the other deletion and thus able to amplify the shorter segment when alone or mixed with full length DNA.

To determine how well the amplicons generated by these primers performed in NGS and to assess reproducibility, we sequenced DNA from seven families, the NIST forensic standards (29), and other individuals. The NIST samples were originally characterized by Sanger sequencing but they were subsequently resequenced using NGS in order to better detect heteroplasmy (30). For the NIST and all other samples discussed below, we have ignored some of the hypervariable regions because they have no impact on disease. Following the lead of others (25), positions 302–306, 513–526, 566–573, and 16181–16194 are not included in the analyses below. For the NIST samples, there is an exact match of our results with both Sanger and NGS for homoplasmic variants. With heteroplasmic variants, the previously reported frequency range for the C64T variant in SRM-CHR was 30–33% for six replicates and we found 29.6%. The previously reported SRM-9947A frequency ranges for G1393A, G3242A, and T7861C were 15–18%, 3–5%, and 68–89%. We found frequencies within these ranges (G1393A: 15.8%, G3242A: 3.3%, T7861C: 88%). This excellent agreement with well-characterized samples encouraged us to examine other samples.

One family group consisted of 12 related HapMap samples with a maternal inheritance set that included the mother (NA12878) and 11 children who should, if there were no mutations, all have identical sequences. A second family was chosen on the basis of longevity and consisted of two nonagenarian siblings (A1 and A2) whose mother was a centenarian, three children (B1, B2, B3) from a nonagenarian daughter and three grandchildren (C1, C2, C3). Pedigrees for the HapMap and longevity families are shown in Fig 3 and 4. Other families with known mitochondrial disease are described below.

**Fig 3.**
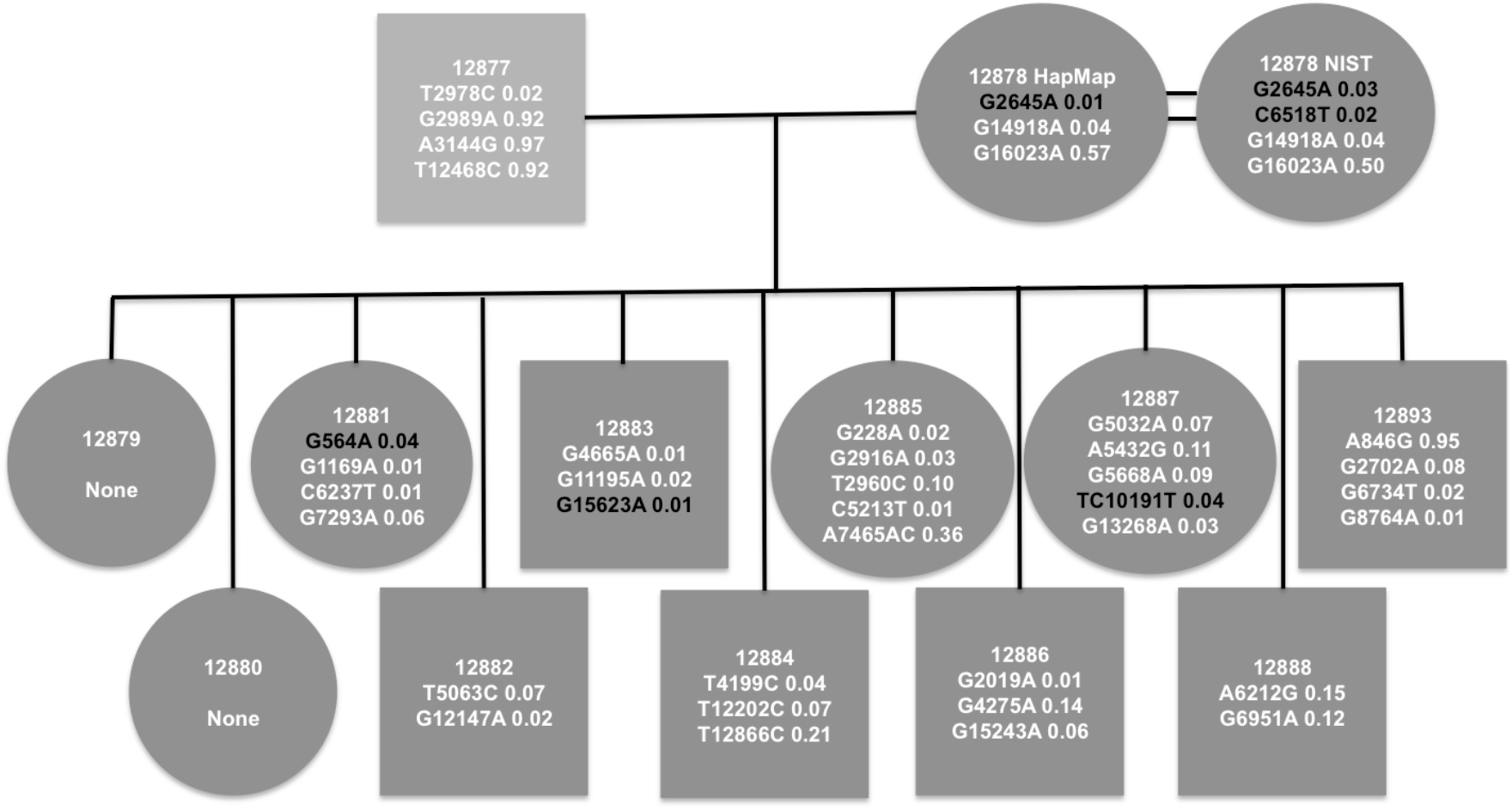
HapMap family pedigree. NA12878 family’s mtDNA is in the H13a1a1a haplogroup (43) with variants C2259T, A4745G, G7337A, T13326C, C13680T, G14831A, and C14872T. Each heteroplasmy in a maternal line is shown with frequency. Those in white were observed previously (31) while those in black were not. No shared heteroplasmy was observed in this family.

**Fig 4.**
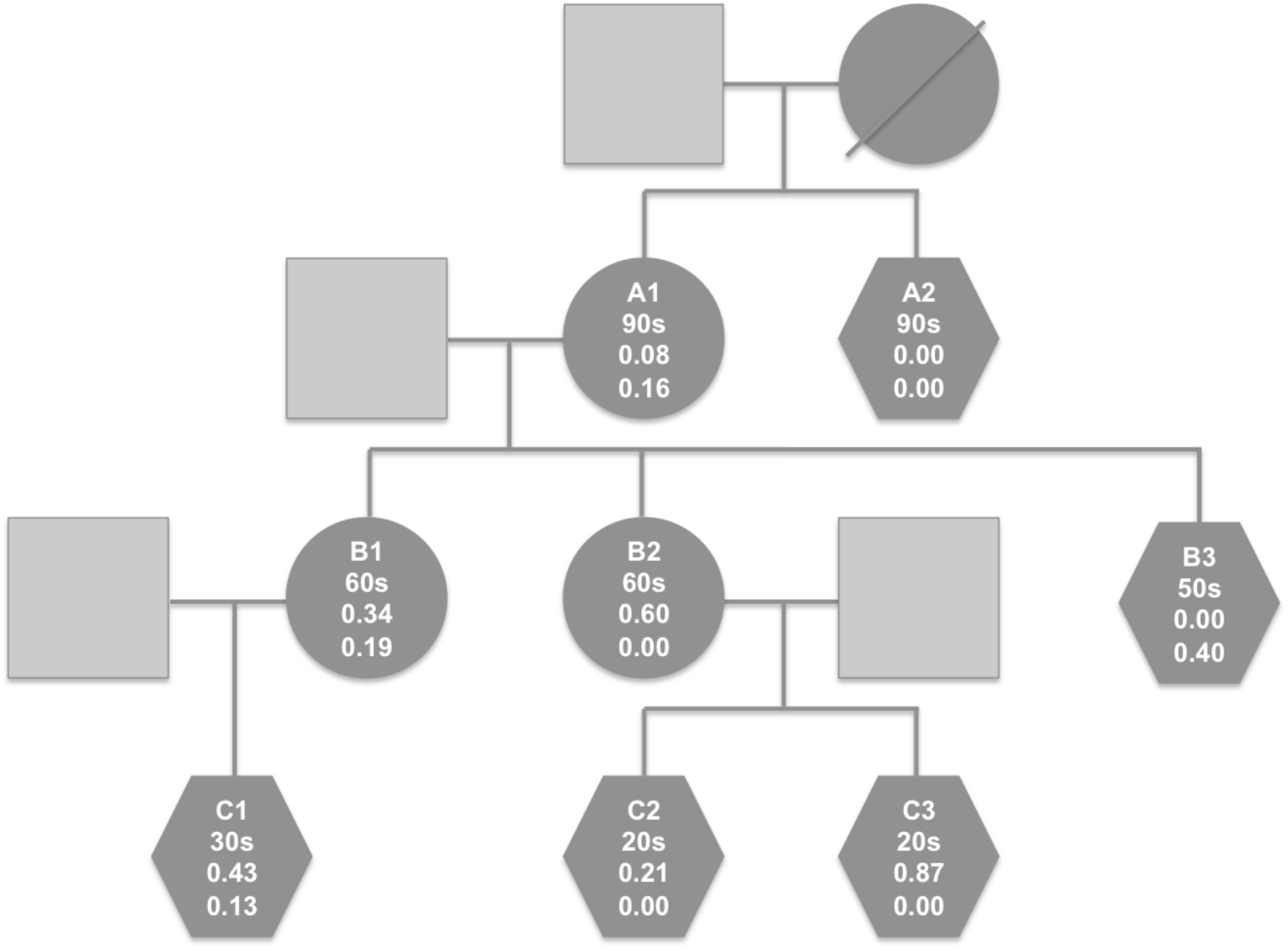
Longevity family pedigree. The numbers in each shape represent the individual ID, the age at which the sample was taken, the frequency of T3290C, and the frequency of T16075C. This family’s DNA is in the T2e haplogroup (43) including variants C150T, T152C, G709A, G1888A, T4216C, A4917G, G8697A, G9948A, T10463C, A11251G, A11812G, G13368A, A14233G, G14905A, C15452A, A15607G, G15928A, T16126C, G16153A, C16294T, and C16296T. Additional variants not used for this haplogroup definition were A73G, C3450T, C3576T, C7028T, G11719A, C14766T, and T16519C.

All the mother-child homoplasmic variants showed the expected inheritance in the HapMap family. However, as shown in Fig 3, 10 of 12 individuals in this family had 1–5 heteroplasmic sites but not a single heteroplasmy was shared among individuals. Of the 38 distinct heteroplasmies, 12 were found at >10% frequency including three at >90% so these are not simply low frequency false positives. Supporting our findings, 34 of these 38 heteroplasmies were observed at similar frequencies in another study using different amplification methods (Supplementary Table 2) (31). Also surprisingly, over half the changes were G->A even though G is strongly underrepresented relative to other bases (~13%) on the reference strand. Despite the preponderance of G->A changes, 3 of the 4 most dramatic frequency changes are A->G (A3144G, A846G, and A6266G). Eight of the 11 HapMap maternal transmissions observed here have at least one site with >5% heteroplasmy and 6 have at least one site with >10% heteroplasmy. All of the heteroplasmic changes observed in the children are rare in the general population with only one having a frequency of >0.4%. The HapMap DNA sequenced is from immortalized cells so it is possible the high frequency of *de novo* heteroplasmies could have arisen in cells during immortalization or culture processes rather than occurring in the original donors. To test this, an independent source (NIST) of NA12878 was examined. The three heteroplasmies found in the original Coriell sample were found in the NIST sample with two of them at slightly different frequencies (Fig 3). In addition, a novel heteroplasmic site was observed, C6518T, at 2% frequency.

The second familial set was sequenced from saliva-derived DNA so potential cell culture artifacts do not limit its interpretation. This family was selected based on its longevity, availability of multiple generations, and the observation of two heteroplasmic sites, T3290C and T16075C, in one of two nonagenarian siblings (A1). The centenarian mother of the nonagenarian siblings died before her DNA could be obtained. The second sibling, A2, had neither heteroplasmic site but was identical to the other sibling at all other mitochondrial positions (T2e haplogroup). Three A1 children, B1, B2, and B3, were also sequenced. B1 had the same two heteroplasmic sites with T16075C found at a very similar frequency as her mother (Fig 4) while T3290C had increased frequency. Both heteroplasmic site frequencies in B1’s child (C1) were similar to the mother. B2, in contrast, had even higher T3290C than the mother, A1, while the T16075C heteroplasmy had disappeared. T16075C was also absent in both of B2’s children while T3290C frequency was still high and drifted in opposite directions in the two children. The third child, B3, lost the other heteroplasmy, T3290C, while T16075C increased in frequency. This multigenerational study shows the dynamic nature of heteroplasmy variation.

The different frequencies of the two heteroplasmies in the longevity family and the fact that both disappear independently suggested that they might reside on separate mtDNA molecules. Short read sequencing is unable to confirm this. Use of a long read technology like Oxford Nanopore has been used previously with mtDNA to assess contamination issues but the higher intrinsic error rate limited its ability to detect low frequency mutations (32). However, once heteroplasmy frequencies are known via short-read sequencing, it may be possible to determine whether variants are on the same molecule as long as they can be amplified on the same fragment. Because the two heteroplasmies in this family are located on different fragments with our standard long range PCRs, alternative primers were chosen to provide a single fragment containing both sites. After amplification and ligation of barcodes according to Oxford Nanopore protocols, the resultant DNA was sequenced using a Minion. To minimize potential artifacts, only barcoded reads were examined. The relevant sites were identified by searching for exact matches to 8 and 9 nt segments with the variant in the middle of the sequence (TCAAYTCCTC for T3290C and CCCAYCAAC for T16075C and their reverse complements). To be included, a read had to contain both sequences separated by 3785 +/−100 bp with the sequences in the correct orientation. Of the 302 sequence reads meeting these criteria for B1, 158 were reference at both sites, 87 were 3290 variant/16075 reference, 38 were 3290 reference/16075 variant, and 19 were variant at both sites. The T3290C allele frequency of 35.1% matched the previously determined frequency (using different primers and sequencing system) of 34%. Similarly, the T16075C frequency was 18.9% using nanopore sequencing compared to 19% previously found. Based on the overall variant frequencies, one would expect 6.6% of molecules to contain variants at both sites if the variants were randomly located. We found 6.3% of the molecules contained both variants, indicating they were indeed random. Similar results were found with C1 though the allele frequencies did not match perfectly (43 vs 50% for T3290C and 13 vs 9% for T16075C). The double mutant molecules were found 3.6% of the time rather than the expected 4.7%.

Because sample preparation for the Oxford Nanopore required both amplification to generate sufficient sample amounts and ligation for barcoding, a control sample was tested to ensure the findings were not artifactual. HapMap samples NA12878 and NA24631 were mixed and subjected to the same amplification and ligation conditions as the dual heteroplasmy samples. These samples contain numerous sites that differ from each other and 10 were chosen for analysis, 4811, 3970, 2706, 1824, 1005, 16304, 16051, 14872, 14831, and 13928. Four bases on each side of the variant site were used to identify sequence locations. Molecules with a barcode and at least 6 identifiable sites in the correct order on the same strand were examined. Sites had to be approximately the right distance from each other (offset distance allowed varied by sites). Molecules were then examined to see if they exclusively matched NA12878 or NA24631 or a mix of the two. Chimeric molecules could arise due to any of many potential technical artifacts such as template switching during amplification, improper ligation during barcoding, or sequencing read errors. While 79% molecules did indeed match only one of the HapMap samples, the remaining molecules did not. Of those, 15% contained a single mismatch site that could have arisen from a sequencing error or other artifact. Most of the singly modified sites were at one of the terminal sites so it is also possible these arose from spurious ligation events. 5% of the molecules contained 2 or more incorrect positions and these were always in blocks, suggesting template switching during amplification or ligation artifacts. As a result, the observation that heteroplasmic sites are randomly assorted on mtDNA molecules must be viewed cautiously. Examples of alignments of three reads with correct variant sequences throughout their entire lengths are shown in Supplementary File 2. In the putative 8056 bp reads (range 7983–8456), the sequence is 82.6–87.1% correct with additional gap errors of 7.9–9.6%. Blast alignments for three reads are provided in Supplementary materials.

We also examined five families shown to have mitochondrial disorders via Sanger sequencing of the proband (Table 3). One patient was confirmed to have a heteroplasmic pathogenic variant, G8969A, using the bait-capture methodology. At the pathogenic site, coverage was 265x with 83.8% of the reads corresponding to the pathogenic 8969A. When this sample was subjected to long range PCR, independent amplicons yielded total coverage of 9545x and 10,828x. 8969A reads were found at 80.6% and 81.9%. With the bait-captured sample, the low coverage made it impossible to determine whether other non-reference bases were elevated relative to background. With the lrPCR samples, the T signal is 2.7% in both samples which is much higher than the background signal of 0.2–0.4% in other samples. The proband’s mother had 1.9% and 1.8% 8969A with coverage of 9359x and 10,862x. Background 8969A signal in other individuals is ~0.1%, suggesting the mother’s heteroplasmy at this position is low but real. Whether the blood heteroplasmy reflects heteroplasmy in other tissues is unknown. The father and a sibling had background levels of 8969A (0.05–0.16%). In two other, unrelated families with the same mutation, the probands had 99.2% and 81.2% 8969A. The two mothers had lower but easily detectable heteroplasmy (12.6% and 37.1%) and two unaffected siblings had undetectable heteroplasmy (<0.1%). While the NGS results were clear, Sanger sequencing had been unable to resolve these differences even when the mutation site was known.

**Table 3:** Base frequencies at position 8969 in families with a mitochondrial disorder

| Freq at 8969 | Long Range PCR 1 |  |  |  | Long Range PCR 2 |  |  |  | Hybridization Bait Capture |  |  |  |
| --- | --- | --- | --- | --- | --- | --- | --- | --- | --- | --- | --- | --- |
| Individual | G | A | T | C | G | A | T | C | G | A | T | C |
| Proband 1 | 0.62 | 99.19 | 0.05 | 0.14 | 0.44 | 99.21 | 0.04 | 0.32 |  |  |  |  |
| Mother 1 | 87.24 | 12.63 | 0.04 | 0.09 | 87.27 | 12.59 | 0.08 | 0.06 |  |  |  |  |
| Proband 2 | 16.18 | 80.56 | 0.52 | 2.73 | 14.85 | 81.91 | 0.58 | 2.65 | 12.83 | 83.77 | 1.13 | 2.26 |
| Mother 2 | 96.86 | 1.94 | 0.92 | 0.27 | 97.02 | 1.80 | 0.82 | 0.35 |  |  |  |  |
| Sibling 2 | 98.76 | 0.16 | 0.84 | 0.23 | 98.94 | 0.05 | 0.85 | 0.17 | 97.10 | 0.36 | 1.09 | 1.45 |
| Father 2 | 99.77 | 0.12 | 0.07 | 0.05 | 99.75 | 0.08 | 0.14 | 0.03 |  |  |  |  |
| Proband 3 | 17.80 | 81.88 | 0.04 | 0.26 | 17.55 | 82.27 | 0.05 | 0.13 |  |  |  |  |
| Mother 3 | 62.82 | 36.88 | 0.16 | 0.13 | 62.23 | 37.51 | 0.13 | 0.13 |  |  |  |  |
| Sibling 3A | 99.77 | 0.06 | 0.12 | 0.05 | 99.80 | 0.02 | 0.10 | 0.06 |  |  |  |  |
| Sibling 3B | 99.79 | 0.09 | 0.07 | 0.03 | 99.78 | 0.08 | 0.11 | 0.02 |  |  |  |  |
| Father 3 | 99.75 | 0.13 | 0.08 | 0.04 | 99.72 | 0.18 | 0.08 | 0.02 |  |  |  |  |
| Proband 4 | 98.70 | 0.08 | 0.97 | 0.25 | 98.65 | 0.12 | 0.93 | 0.30 | 97.54 | 0.00 | 1.43 | 1.02 |
| Mother 4 | 98.80 | 0.09 | 0.98 | 0.13 | 98.92 | 0.10 | 0.80 | 0.17 | 97.18 | 0.00 | 1.69 | 1.13 |
| Father 4 | 98.87 | 0.15 | 0.81 | 0.17 | 98.86 | 0.11 | 0.93 | 0.10 | 95.96 | 0.00 | 2.53 | 1.52 |
| Proband 5 | 98.77 | 0.09 | 0.79 | 0.35 | 98.79 | 0.10 | 0.89 | 0.20 | 93.75 | 1.56 | 3.13 | 1.56 |
| Mother 5 | 98.98 | 0.07 | 0.77 | 0.18 | 98.87 | 0.11 | 0.84 | 0.18 | 95.59 | 0.00 | 1.47 | 2.94 |

Heteroplasmy rates for SNVs are obtained straightforwardly from the relative read numbers. It is not as simple for deletions because different sized fragments can amplify at different rates, causing issues with accurate quantitation. This is easily seen in Fig 1 where the deletion fragments predominate even when the full-length DNA is in excess. We have used literature qPCR methods (33) to measure the relative copy number of the deletions relative to the backbone DNA. In order to use the appropriate qPCR primers, it is necessary to localize the deletions with sequencing first. With the two deletion samples we had access to, qPCR methods using triplicate measurements indicated the common deletion was present at 86.5% while the 10/15 deletion was present in 82.2% of molecules. Because this method requires more DNA as well as a separate workflow and instrument compared to NGS sequencing, we sought to identify another method for measuring deletions that did not require qPCR.

To circumvent the amplification bias, we mixed nominally equal amounts of DNA from HapMap sample NA24631 with DNA from patients with the heteroplasmic, large deletions described above. Even this is not straightforward as equivalent amounts of total DNA may contain different proportions of mtDNA. However, an accurate assessment of the relative mtDNA concentrations can be obtained by examining variant frequencies in Fragment 2, which is not affected by the deletions. This fragment is identically sized in both samples so should amplify identically with both mtDNAs. As shown in Table 4, combining equal amounts of NA24631 with either the Common deletion DNA or the 10/15 deletion DNA indicated the mixes actually contained 73.9% and 54.6% NA24631 mtDNA, respectively, based on variant frequency in Fragment 2 (positions 16510–7891). Because these measurements are of mixtures, the variant frequency measurement is not heteroplasmy but rather an indication of how much DNA from each component is present in the mixture.

**Table 4:** Heteroplasmy in Deletion Mixtures. The two deletion samples (Haplogroups H11a and L2c2b2) were mixed with HapMap sample NA24631 (F2b1). Each mixture was then amplified using primers 1/2 and 3/4 and sequenced. Because there were many differences in sequence between the mtDNAs, there were many apparent heteroplasmies. These were averaged over 3 regions: all of Fragment 2 and two segments of Fragment 1, both inside and outside the deletion region. This allows heteroplasmy of the original deletion sample to be determined. The overlapping segments of Fragments 1 and 2 were omitted from the analysis. The heteroplasmic site C12969T in NA24631 was not used in the analysis. G10310A was omitted from the 10/15 analysis because it was a significant outlier that could have been impacted by alignment issues near the deletion breakpoint. For many variant sites, the test sample varied from the reference and NA24631 had the reference base. For consistency in fractions, these sites were treated as if NA24631 was non-reference even though it actually matched the reference sequence.

| Deletion | Mixing Ratio | Location | Variant Frequency on Deleted | Variant Frequency on NA24631 | Number of SNVs |
| --- | --- | --- | --- | --- | --- |
| 10/15 | 50:50 | Fragment 2 | 0.454 | 0.546 | 15 |
| 10/15 | 50:50 | Fragment 1 Outside del | 0.877 | 0.123 | 8 |
| 10/15 | 50:50 | Fragment 1 Inside del | 0.179 | 0.821 | 10 |
| 10/15 | 25:75 | Fragment 2 | 0.199 | 0.801 | 15 |
| 10/15 | 25:75 | Fragment 1 Outside del | 0.678 | 0.322 | 8 |
| 10/15 | 25:75 | Fragment 1 Inside del | 0.065 | 0.935 | 10 |
| Common | 50:50 | Fragment 2 | 0.261 | 0.739 | 34 |
| Common | 50:50 | Fragment 1 Outside del | 0.829 | 0.171 | 17 |
| Common | 50:50 | Fragment 1 Inside del | 0.052 | 0.948 | 19 |

In order to determine the true heteroplasmy of deletions in the patient samples, Fragment 1 (spanning the latter half of the circular genome, positions 7709–16569 and 1–5) was separated into two regions for analysis: 1) the DNA segments unaffected by the deletion; 2) the DNA segments present only in full-length DNA and not in the deletions. Plotting of variant frequency versus position in mtDNA is shown in Supplementary Fig 2 where the regions can be clearly distinguished based on changes in variant frequency. The amplified DNA mixture contains deleted test DNA, full-length test DNA, and NA24631 with all three DNAs contributing to the Fragment 2 measurement but only the latter two DNAs to the Fragment 1/inside del measurement. Neither of these measurements is affected by amplification efficiency as only the same size DNA fragments are compared (in contrast to the measurement of Fragment 1 outside the deletion region where different size DNA fragments contribute). The fractional contribution of all three DNAs (deleted test DNA, full-length test DNA, and NA24631) can be calculated because the relative amounts of the DNAs are constant and the fractions in each measurement must add to 1.

The variant frequency of NA24631 is higher in the deleted region, increasing from 0.546 to 0.821 (10/15) and from 0.739 to 0.948 (Common), reflecting the lost contribution from the deleted test DNA in that measurement. The full-length test DNA fraction in the Fragment 2/Inside del measurement increases by the same proportion as the HapMap DNA, so the variant frequency of 0.179 (10/15) and 0.065 (Common) can be used to calculate the true fractional contributions: 0.119 in 10/15 and 0.041 in Common. See Supplementary Table 1 for details on how the calculations are performed. With these values for the fractional contribution of the full-length test sample and the previously determined fraction of NA24631, the fractional contribution of deleted test sample can be calculated as 0.335 (10/15) and 0.220 (Common). This yields overall deletion heteroplasmies of 74% for 10/15 and 84% for the Common deletion. This analysis was repeated for the 10/15 deletion with a 75:25 mix of NA24631:10/15 DNA, resulting in a calculated heteroplasmy of 28% full-length and 72% deleted. This measurement, though close to the value generated using the 50:50 mix, is probably less accurate because the variant frequencies were lower and less accurate. The accuracy of these estimates is enhanced by choosing a spike-in DNA with many sequence differences relative to the test DNA as is the case with NA24631.

Because urine cells provide a useful, potentially independent tissue source of mtDNA, we sought to better understand the tissue origin of these cells to ensure they were not derived from blood like saliva DNA has been shown to be in some cases. RNA was prepared from freshly collected cells suspended in the urine and analyzed via digital gene expression (DGE) with 15–40M reads/sample (34, 35). The most highly expressed genes were ribosomal proteins, translational elongation factors, housekeeping, and similar genes. These genes provide little information about tissue origin because they are expressed in nearly all cells. To focus on tissue-specific RNAs, DGE data was obtained from frozen tissue samples (liver, kidney cortex, kidney medulla and muscle) and other fresh fluids (saliva and whole blood). Data was also obtained from the Ion Torrent community website for the Universal Human Reference (UHR), the Human Brain Reference (HBR), and a Lung FFPE tumor/normal pair. Genes that had the highest ratio of expression in the urine cells compared to the UHR samples are shown in Table 5. Genes are sorted by urine/UHR expression ratio.

**Table 5:**
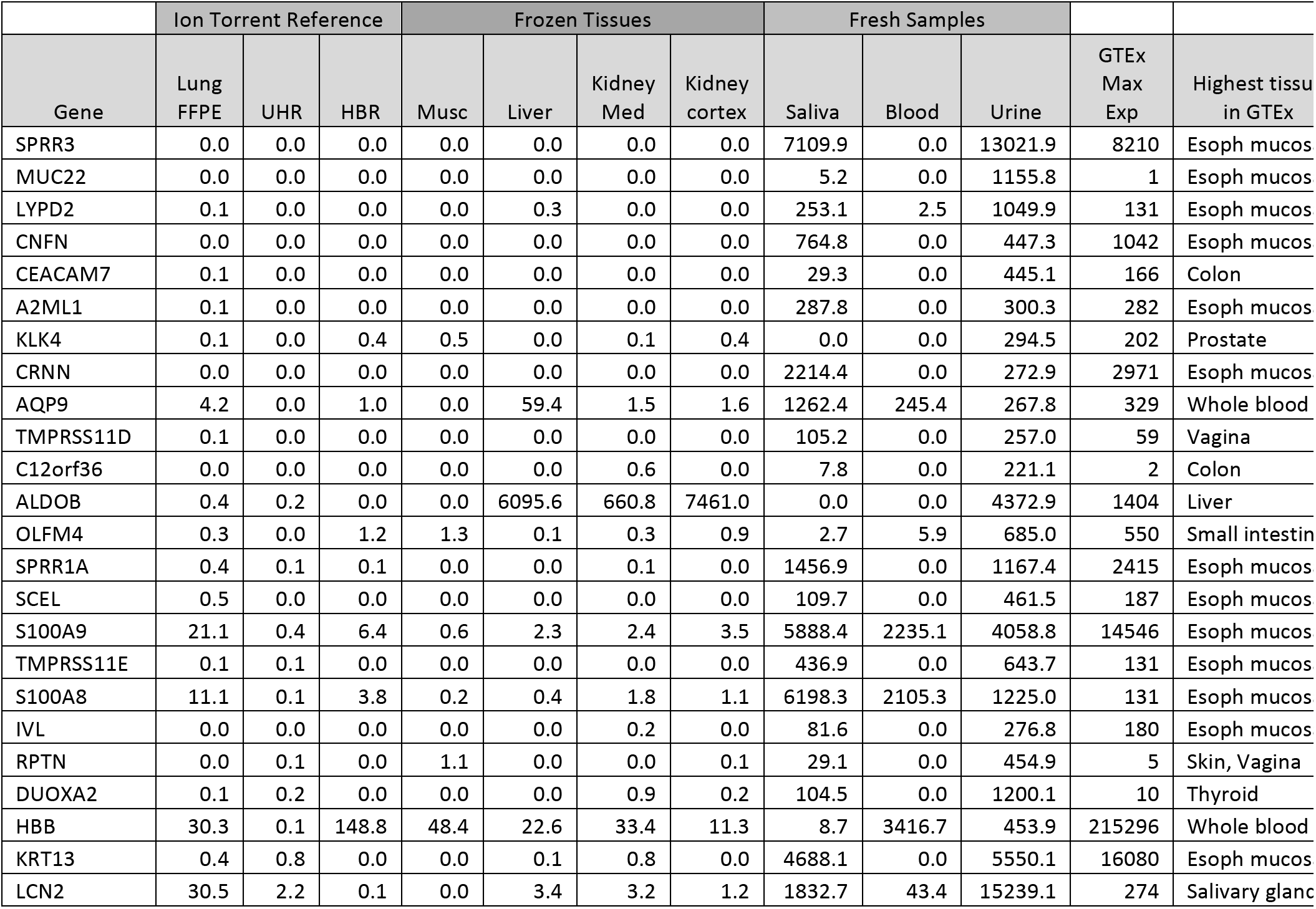
Digital Gene Expression in Urine Cells. Digital Gene Expression was carried out on RNA from urine cells, saliva and blood as well as frozen tissue samples (muscle, liver, kidney medulla and cortex) and compared to the Universal Human Reference (UHR), human brain reference (HBR), and lung cancer samples from the Ion Torrent website. The genes with the highest ratio of urine/UHR reference samples are listed. 0.01 RPM was added to each UHR expression level to allow a ratio to be calculated when UHR expression was 0. Maximum RNA Seq expression values and the highest expressing tissue found in GTEx are provided (RPKM) for comparison. “Esoph mucosa” means mucosal cells from the esophagus. Other mucosal cells were not in the database. Normalization by RNA length to generate RPKM was not performed. See reference (35) for a description of the differences between DGE and RNA Seq expression and the impact of length normalization.

|  | Ion Torrent Reference |  |  | Frozen Tissues |  |  |  | Fresh Samples |  |  |  |  |
| --- | --- | --- | --- | --- | --- | --- | --- | --- | --- | --- | --- | --- |
| Gene | Lung<br>FFPE | UHR | HBR | Musc | Liver | Kidney<br>Med | Kidney<br>cortex | Saliva | Blood | Urine | GTEX<br>Max<br>Exp | Highest tissu<br>in GTEX |
| SPRR3 | 0.0 | 0.0 | 0.0 | 0.0 | 0.0 | 0.0 | 0.0 | 7109.9 | 0.0 | 13021.9 | 8210 | Esoph mucos |
| MUC22 | 0.0 | 0.0 | 0.0 | 0.0 | 0.0 | 0.0 | 0.0 | 5.2 | 0.0 | 1155.8 | 1 | Esoph mucos |
| LYPD2 | 0.1 | 0.0 | 0.0 | 0.0 | 0.3 | 0.0 | 0.0 | 253.1 | 2.5 | 1049.9 | 131 | Esoph mucos |
| CNFN | 0.0 | 0.0 | 0.0 | 0.0 | 0.0 | 0.0 | 0.0 | 764.8 | 0.0 | 447.3 | 1042 | Esoph mucos |
| CEACAM7 | 0.1 | 0.0 | 0.0 | 0.0 | 0.0 | 0.0 | 0.0 | 29.3 | 0.0 | 445.1 | 166 | Colon |
| A2ML1 | 0.1 | 0.0 | 0.0 | 0.0 | 0.0 | 0.0 | 0.0 | 287.8 | 0.0 | 300.3 | 282 | Esoph mucos |
| KLK4 | 0.1 | 0.0 | 0.4 | 0.5 | 0.0 | 0.1 | 0.4 | 0.0 | 0.0 | 294.5 | 202 | Prostate |
| CRNN | 0.0 | 0.0 | 0.0 | 0.0 | 0.0 | 0.0 | 0.0 | 2214.4 | 0.0 | 272.9 | 2971 | Esoph mucos |
| AQP9 | 4.2 | 0.0 | 1.0 | 0.0 | 59.4 | 1.5 | 1.6 | 1262.4 | 245.4 | 267.8 | 329 | Whole blood |
| TMPRSS11D | 0.1 | 0.0 | 0.0 | 0.0 | 0.0 | 0.0 | 0.0 | 105.2 | 0.0 | 257.0 | 59 | Vagina |
| C12orf36 | 0.0 | 0.0 | 0.0 | 0.0 | 0.0 | 0.6 | 0.0 | 7.8 | 0.0 | 221.1 | 2 | Colon |
| ALDOB | 0.4 | 0.2 | 0.0 | 0.0 | 6095.6 | 660.8 | 7461.0 | 0.0 | 0.0 | 4372.9 | 1404 | Liver |
| OLFM4 | 0.3 | 0.0 | 1.2 | 1.3 | 0.1 | 0.3 | 0.9 | 2.7 | 5.9 | 685.0 | 550 | Small intestin |
| SPRR1A | 0.4 | 0.1 | 0.1 | 0.0 | 0.0 | 0.1 | 0.0 | 1456.9 | 0.0 | 1167.4 | 2415 | Esoph mucos |
| SCEL | 0.5 | 0.0 | 0.0 | 0.0 | 0.0 | 0.0 | 0.0 | 109.7 | 0.0 | 461.5 | 187 | Esoph mucos |
| S100A9 | 21.1 | 0.4 | 6.4 | 0.6 | 2.3 | 2.4 | 3.5 | 5888.4 | 2235.1 | 4058.8 | 14546 | Esoph mucos |
| TMPRSS11E | 0.1 | 0.1 | 0.0 | 0.0 | 0.0 | 0.0 | 0.0 | 436.9 | 0.0 | 643.7 | 131 | Esoph mucos |
| S100A8 | 11.1 | 0.1 | 3.8 | 0.2 | 0.4 | 1.8 | 1.1 | 6198.3 | 2105.3 | 1225.0 | 131 | Esoph mucos |
| IVL | 0.0 | 0.0 | 0.0 | 0.0 | 0.0 | 0.2 | 0.0 | 81.6 | 0.0 | 276.8 | 180 | Esoph mucos |
| RPTN | 0.0 | 0.1 | 0.0 | 1.1 | 0.0 | 0.0 | 0.1 | 29.1 | 0.0 | 454.9 | 5 | Skin, Vagina |
| DUOXA2 | 0.1 | 0.2 | 0.0 | 0.0 | 0.0 | 0.9 | 0.2 | 104.5 | 0.0 | 1200.1 | 10 | Thyroid |
| HBB | 30.3 | 0.1 | 148.8 | 48.4 | 22.6 | 33.4 | 11.3 | 8.7 | 3416.7 | 453.9 | 215296 | Whole blood |
| KRT13 | 0.4 | 0.8 | 0.0 | 0.0 | 0.1 | 0.8 | 0.0 | 4688.1 | 0.0 | 5550.1 | 16080 | Esoph mucos |
| LCN2 | 30.5 | 2.2 | 0.1 | 0.0 | 3.4 | 3.2 | 1.2 | 1832.7 | 43.4 | 15239.1 | 274 | Salivary glanc |

The expression profiles of the listed urine-selective genes were compared to the expression levels in 51 tissues found in the GTEx database (gtexportal.org/). Because the GTEx data was generated using classical RNA Seq and our data was generated using DGE that is independent of transcript length, the normalized units differ, RPKM versus RPM. 5 of the top 6 and 12 of the top 20 most selectively expressed genes in urine cells had their highest GTEx expression levels in esophageal mucosa cells. MUC22, DUOXA2, and C12orf36 are expressed more than 100x higher and RPTN, LCN2, CXCL6, GABRP, DUOX2, MT1H, GPR110, MT1L, PRSS22, GBP5, SPNS2, and TMPRSS2 are expressed 20–100x higher in urine cells than any tissue reported in GTEx. The GTEx database does not include all tissues, but the comparison suggests that most cells in the urine are probably mucosal cells that slough off into the urine. Other cell types must contribute RNA/DNA to a lesser extent.

Similarities in tissue gene expression patterns were examined in two ways. To look at overall similarities, all genes with >1 RPM in at least one tissue were compared pairwise across all tissues (Table 6A). Not unexpectedly, the highest correlation was kidney medulla versus kidney cortex with r=0.991 while the least similar pair was muscle versus kidney medulla (0.048). The UHR sample, intended to be representative of all tissues, did indeed have the smallest range of r values (0.3390.716). Saliva, blood, and urine had modest similarity (0.510–0.608).

**Table 6:** Gene expression similarities across tissues. A) Pearson’s r correlation between tissue expression levels (omitting genes expressed at less than 1 RPM in these samples) B) Sharing of tissue specific genes Genes with either >100x (top right) or >10x (lower left) greater expression in a tissue relative to UHR were compared between tissues. The number of such genes (10x/100x) is shown in the diagonal for each tissue and the number shared with the other listed tissues is shown above (>100x) or below (>10x) the diagonal.

|  | UHR | HBR | Urine | Saliva | Blood | Muscle | Liver | Kidney Cortex | Kidney Medulla |
| --- | --- | --- | --- | --- | --- | --- | --- | --- | --- |
| FFPE Lung | 0.783 | 0.378 | 0.485 | 0.434 | 0.527 | 0.211 | 0.653 | 0.619 | 0.646 |
| UHR |  | 0.380 | 0.561 | 0.415 | 0.716 | 0.339 | 0.396 | 0.360 | 0.362 |
| HBR |  |  | 0.225 | 0.229 | 0.250 | 0.118 | 0.236 | 0.247 | 0.252 |
| Urine |  |  |  | 0.580 | 0.607 | 0.214 | 0.350 | 0.373 | 0.350 |
| Saliva |  |  |  |  | 0.510 | 0.122 | 0.282 | 0.265 | 0.271 |
| Blood |  |  |  |  |  | 0.262 | 0.184 | 0.175 | 0.161 |
| Muscle |  |  |  |  |  |  | 0.060 | 0.052 | 0.048 |
| Liver |  |  |  |  |  |  |  | 0.814 | 0.831 |
| Kidney Cortex |  |  |  |  |  |  |  |  | 0.991 |

| 10x v 100x > | FFPE Lung | HBR | Urine | Saliva | Blood | Muscle | Liver | Kidney Cortex | Kidney Medulla |
| --- | --- | --- | --- | --- | --- | --- | --- | --- | --- |
| FFPE Lung | 412/19 | 3 | 3 | 4 | 4 | 0 | 0 | 0 | 0 |
| HBR | 145 | 1031/100 | 1 | 8 | 1 | 3 | 0 | 1 | 1 |
| Urine | 65 | 53 | 585/110 | 56 | 21 | 0 | 4 | 6 | 3 |
| Saliva | 62 | 33 | 299 | 755/149 | 50 | 0 | 1 | 2 | 0 |
| Blood | 63 | 35 | 167 | 348 | 564/81 | 0 | 0 | 0 | 0 |
| Muscle | 44 | 72 | 31 | 31 | 27 | 423/105 | 1 | 1 | 1 |
| Liver | 45 | 42 | 91 | 55 | 43 | 42 | 513/82 | 5 | 2 |
| Kidney Cortex | 42 | 61 | 126 | 32 | 26 | 31 | 123 | 489/51 | 16 |
| Kidney Medulla | 58 | 75 | 128 | 50 | 44 | 46 | 121 | 344 | 633/28 |

Rather than looking at >18K genes with expression levels > 1 RPM in any tissue, it is also instructive to look at only the tissue specific genes (Table 6B). Genes with either >100x or >10x expression in each tissue relative to UHR were compared. Genes with >100x expression ranged from a low of 19 with lung FFPE to 149 for saliva.

Similarly, there was a low of 412 genes with >10x expression in lung FFPE ranging to a high of 1031 in HBR. The highest degree of sharing of >100x genes was between urine and saliva (56 out of a possible 110) and the highest degree of sharing of >10x genes was between blood and saliva (348 out of a possible 564). These comparisons show that there are similarities among the different fluids evaluated but there are also sufficient differences to indicate that blood cells do not contribute substantially to saliva and urine signals in these healthy individuals.

To confirm that sequencing of urine mtDNA yielded the same result as saliva DNA, multiple samples were sequenced including one of the samples with heteroplasmies. Because of the issues with effective amplification/sequencing of urine mtDNA due to its low yield, low-level heteroplasmies (up to 3%) were ignored. The single heteroplasmic sample for which both saliva and urine were available yielded very similar frequencies comparing the two (T3290C: 0.34 vs 0.37; T16075C: 0.19 vs 0.23) and homoplasmic variants were identical.

## Discussion

When DNA sequencing was restricted to Sanger methodology, the ability to sequence the whole mitochondrial genome and detect low-level heteroplasmies was very limited. The NIST forensic standards demonstrate this clearly where NGS analysis identified two heteroplasmies in SRM-9947A that were missed by Sanger sequencing (30). Despite this, Sanger sequencing of mtDNA could still be used effectively for forensic and genealogical purposes because these applications do not generally require information from the whole mitochondrial genome. However, a complete knowledge of mtDNA is required in order to detect potential causal variants in disease. While the advent of NGS technologies opened up mtDNA to a more complete analysis, technical aspects of mtDNA sequencing remain more challenging than with nuclear DNA and their impact needs to be understood in order to obtain results accurate and reproducible enough for clinical diagnostics. For example, knowing that heteroplasmy can vary significantly across both tissues and generations affects how one chooses which samples to examine and how to sequence them. In order to reliably achieve detection of heteroplasmies down to 1% or less, care must be taken to choose sample preparation methods and enzymes that are highly accurate or the background error rate will obscure true heteroplasmies. The appearance of novel heteroplasmic sites in most of HapMap samples we sequenced raised concerns about the origin of the variants. 34 of the 38 sites we identified were found by others (31) (see Supplementary Table 2 for frequency comparison) using different amplification methods. Novel heteroplasmies have been similarly observed in other HapMap samples (36, 37). With access to DNA only from immortalized cells for the HapMap samples, it is impossible to examine tissue-specific heteroplasmies or even to rule out whether the new heteroplasmies occurred naturally. Their high frequency makes it highly likely that they were induced during the immortalization/culture process. This hypothesis was supported by our sequencing of two different NA12878 DNA preparations. Though all homoplasmic and most heteroplasmic sites were identical between DNAs, there were three differences. One new heteroplasmic site was found in the NIST DNA and two other sites changed frequencies somewhat. The “novel” site may, in fact, have been present in the original DNA but more highly selected in the second DNA preparation or it may have arisen later.

The highly asymmetric nature of the novel HapMap heteroplasmies, 22 of 38 being G->A despite only 13% of bases on the reference strand being G, suggests a nonrandom mechanism for the generation/selection of these variants. Others have examined changes in mtDNA in tumors with similar results (38). 28 null variants were fixed in tumors with lower activity seemingly selected for. 18 of these were frameshifts but 9 of the 10 SNVs were G>A. The observation that 3 of the 4 most prevalent HapMap heteroplasmies are among the 6 A->G variants is also suggestive of non-random effects on replication and repair but the mechanistic basis for both these phenomena is beyond the scope of this work.

Like the cell-based HapMap samples, the two NIST standard mtDNAs that we examined are prepared from immortalized cell lines. They also have heteroplasmies that are rarely found in the general population and 2 of the 4 were G->A changes though the two with the highest frequency were not G->A. These samples provide a wide range of heteroplasmic frequencies that permits the sensitivity of various methods to be determined. These standard DNAs were generated from cell lines, and maintaining reliable heteroplasmy standards will require continual monitoring as heteroplasmic frequencies could change in future DNA preparations.

In contrast to the HapMap samples, 14 maternal mtDNA transmissions among seven other families that we tested resulted in the detection of no *de novo* heteroplasmies. In families with pre-existing heteroplasmy, the frequency usually changed, sometimes substantially, but no new sites were observed. There are few literature estimates of the rate of *de novo* heteroplasmy generation among healthy individuals because there are few studies that have examined the whole mtDNA at sufficient depth and accuracy to provide solid numbers. One study with sufficient coverage depth (~20,000x) used the number of heteroplasmies and the rate of heteroplasmic drift among mother/child pairs to allow calculation of *de novo* variants to about 1 per 100 healthy births (39). This rate is consistent with our findings of no *de novo* variants in saliva/blood samples but is not consistent with a natural cause for the high *de novo* rate observed in HapMap samples.

Even with the artifactual heteroplasmies in the HapMap samples, the varied levels are useful for assessing the ability of various protocols to generate reproducible results. In theory, each method should give identical results but each is also subject to its own potential technical artifacts that could lead to discrepancies. Using these sensitive methods, it is possible to follow inherited heteroplasmies across generations and detect it down to 1% or less. The ability to detect such low levels can be important. The mother of proband 2 (Table 3) was observed to have the pathogenic G8969A variant at very low levels so is at risk of further transmissions but Sibling 2 does not have the variant so is not at risk of disease or variant transmission. Both pieces of information are useful in health planning. The heteroplasmies in the longevity family are apparently neutral and thus have no health consequences and their frequencies drift in both directions. Interestingly, this family’s mtDNA is in the T2e haplogroup that has been associated with longevity (40). The two observed heteroplasmic sites are not found in the T2e haplogroup though these variants are found in other non-T haplogroups. The heteroplasmic frequencies of the two variants did not appear correlated and transmission might be independent based on each one being lost in separate events so we expected they might be on different molecules. To address this question, we sequenced the molecules using long-read Oxford Nanopore technology. Unfortunately, the current protocols for sample preparation include amplification to attain enough DNA and ligation to add barcodes. Both of these steps can alter the original DNA and this issue was confirmed as problematic by testing mixtures of very similar DNAs. While most DNAs were unaffected, up to 20% could be impacted and thus obscure the small effects we were trying to observe. We were unable to confirm the relative locations of the heteroplasmies because independent molecules that were mixed and subjected to the same protocol showed evidence of artifactual variant sharing. These reads would be useful for consensus sequence determination but not for establishing haplotype of uncommon variants within a larger background of other molecules.

While calculating heteroplasmy with SNVs is simple, doing the same for large deletion variants is not as straightforward with published NGS methods. Potential amplification biases in large versus small fragments can skew the apparent ratio of the two, making accurate determination challenging. Over-counting a deleted segment occurs often (41). The full-length fragments can be virtually undetectable despite being present in substantial amounts as seen in Fig 1. Simply looking at the number of junction reads versus intact reads or coverage depths could result in incorrect estimations as well. To circumvent this problem, we have developed a novel method requiring only NGS data to more accurately assess the fraction of molecules with deletions. The deleted samples were spiked with a constant amount of DNA (both mitochondrial and nuclear) from a highly divergent HapMap sample to allow separation of the amplification effects from true heteroplasmy. The fragment without a deletion should amplify identically in both samples providing the true ratio of mtDNA concentrations from the two samples. This value can then be used to determine the contribution of each sample to the signal in the deleted region, allowing the true ratio of deleted to full length in the test sample to be determined. The large number of SNV differences between the spiked sample and the test sample enhances the accuracy of these measurements. There were 33 and 70 differences in the two pairs of samples, so the variant frequencies can be averaged for higher accuracy. The qPCR and NGS methods agree reasonably well for heteroplasmy determination in these samples. As can be seen in Supplementary Fig 2, the variant frequencies can also be used to generate a low resolution mapping of the deletion. The magnitude of the amplification bias can also be estimated. Based on the excess heteroplasmy due to the deletion fragments in the amplified fragment outside the deletion region, one can estimate an overrepresentation of ~6.8x for the short Common deletion fragment and ~11x for the short 10/15 deletion fragment using these amplification conditions. The larger effect with 10/15 is expected because its deletion is larger. This method of determining heteroplasmy for large deletions is attractive for use in a clinical setting because it relies only on NGS data without the need for other technologies like qPCR, thus simplifying the workflow.

Most mtDNA assays use either saliva or blood as a source of the DNA. Based on studies with bone marrow donors/recipients, a significant fraction of DNA in saliva could be derived from blood (21). Thus, saliva may not provide a fully independent view of mutations that may be occurring across tissues. Most other relevant tissues are inconvenient or dangerous to sample so we examined the easily accessible cells found in urine for their suitability. As shown in Table 1, mitochondrial content is much higher in urine cells than blood or saliva. To ensure that these cells were not primarily derived from blood, RNA expression studies were carried out to assess the tissue origin of cells in the urine. Many of the top 100 genes expressed in whole blood were expressed poorly in urine including some with expression < 1 RPM (ALAS2, HLA-B, CD53) or 1–10 RPM (DEFA3, NKG7, RHOG, SERPINA1). Nearly all genes expressed at high levels in blood and urine are highly expressed in most other tissues also. The only exceptions to this are S100A8 and S100A9, which are highly expressed in blood and urine cells but not most other tissues. However, both are also highly expressed in esophageal mucosal cells. Though the cells in urine likely come from a variety of sources, the best match in the GTEx database is clearly mucosal cells so these likely contribute the largest fraction of DNA to the analysis. While there may be a minor contribution from blood to the sample, it must be a small fraction of the mix so urine provides the independent source that is desirable for analysis of mtDNA. While the urine cells provide the desired independent source of DNA, this DNA is limited by its very low yield. While it is easy to get micrograms of DNA from blood or saliva, nanograms is the relevant unit for urine DNA. This limits the methods that can be used for sequencing and the higher level of amplification required means low-level heteroplasmy will be more difficult to detect due to higher background noise.

The ability to generate accurate results for clinical diagnostic sequencing is important so that families may use that information in planning and treatment. Efforts to prevent transmission of pathogenic mitochondrial variants have begun with a variety of strategies. Selection of embryos with low levels of pathogenic variants has had limited success thus far (10) and efforts to generate “three parent” children have begun but are being subjected to a high degree of regulatory and ethical scrutiny due to the uncertainties surrounding the process. Families making these difficult choices should be able to rely on accurate genetic information on which to base their decisions. High quality, clinical mtDNA sequence is achievable but care must be taken to avoid artifacts that can compromise the data.

## Materials and Methods

Oligonucleotide primers for sequencing and amplification were obtained from IDT. HapMap DNAs were purchased from Coriell Institute for Medical Research (Camden, NJ) and forensic standard DNAs from NIST (Gaithersburg, MD). Saliva DNA was collected using Oragene Dx kits. Urine DNA was collected using the Zymo Research Urine DNA Extraction kit. Frozen tissue samples (liver, kidney, muscle) were obtained from AMSBIO (Cambridge, MA). DNA was isolated from blood using either an Autogen FlexStar or FlexStar+. Other DNA was prepared manually using an Oragene prepIT-L2P (saliva and urine) or QIAgen QIAamp Mini kit (liver, kidney, muscle). DNA sizing was carried out using an Agilent 2200 TapeStation with the Genomic DNA Screen Tape and Reagents. For bait-based capture, custom kits with varying ratios of mitochondrial/nuclear baits were obtained from Agilent and sequenced on the Illumina NextSeq platform. Most long range PCR was carried out using primers as described in Table 1 with either NEB Q5 polymerase or Takara LA Taq. For Oxford Nanopore sequencing, a different segment of mtDNA was analyzed using primers CTGCTATGATGGATAAGATTGAGAGA (4962–4937) and GCAGGAATACCTTTCCTCACAG (13477–13498). Amplified DNA was purified with AMPure beads prior to sequencing. Libraries were prepared with an Agilent SureSelect QXT for whole genome sequencing kit. All NGS libraries were prepared according to manufacturers’ instructions. Sequencing was carried out on Illumina NextSeqs and Ion Torrent PGMs and Protons according to manufacturers’ instructions as described previously (42). Oxford Nanopore libraries were prepared with input of 1μg of amplified long-range PCR product using R9 Ligation Sequencing 1D Kit with Native Barcoding, omitting the fragmentation and DNA repair steps. These samples were loaded on R9 Spot-on Flo-min107 flowcells. MinKNOW software was used for instrument control and basecalling. Barcodes were de-convoluted using EPI2ME agent. Sequences were searched for specific variant sites using 8 or 9mers.

RNA was extracted from tissues using the Qiagen RNA Mini kit, from blood using the Qiagen QIAamp RNA blood Mini kit, from saliva using the Qiagen RNeasy Micro Kit, and from urine using the Zymo Research Urine RNA Isolation Kit. RNA was prepared for Digital Gene Expression as described (34) and sequenced on an Ion Torrent Proton. qPCR analysis of mitochondrial:nuclear DNA ratios was performed according to Phillips et al. (33).

NextSeq data underwent alignment, cleaning, and variant calling according to GATK best practices. Variant calling for mtDNA was performed with GATK v3.5 HaplotypeCaller with ploidy 25 with no duplicate removal. For Ion Torrent data, read alignment, cleaning, and variant calling was performed using Torrent Suite v4.4.

None of the human DNA samples are identifiable from the data provided. Where possible, HapMap and publically available DNAs have been used. Other samples from our CLIA laboratory have been anonymized and used for the improvement and quality control of our assays. Our CLIA laboratory IRB has been through Boston Children’s Hospital.

## Acknowledgments

The Genotype-Tissue Expression (GTEx) Project was supported by the Common Fund of the Office of the Director of the National Institutes of Health, and by NCI, NHGRI, NHLBI, NIDA, NIMH, and NINDS. The data used for the GTEx analyses described in this manuscript were obtained from: the GTEx Portal on 07/24/17. Partial funding for this work was obtained via NIH grants R24 DK099808-05 and R01 DK087992.

